# Co-mutation and exclusion analysis in human tumors, a means for cancer biology studies and treatment design

**DOI:** 10.1101/182501

**Authors:** Soledad Ochoa, Elizabeth Martínez-Pérez, Diego Javier Zea, Miguel Angel Molina-Vila, Cristina Marino-Buslje

## Abstract

**Background:** Malignant tumors originate from genomic and epigenomic alterations, which lead to loss of control of the cellular circuitry. These alterations relate with each other in patterns of mutual exclusion and co-occurrence that affect prognosis and treatment response and highlight the need for multitargeted therapy. However, to the best of our knowledge, there are no systematic reports in the literature of co-dependent and mutually exclusive mutations across all types of cancer. In addition, the studies reported so far generally deal with whole genes instead of specific mutations, ignoring the fact that different alterations in the same gene can have widely different effects.

**Results:** Here we present a systematic analysis of co-dependencies of somatic mutations across all cancer types. Combining multi testing with conditional and expected mutational probabilities, we have found pairs and networks of co-mutations and exclusions, some of them in particular types of cancer and others widespread. We have also determined that driver loci are present in more types of cancer than non driver loci, that they tend to pair within a single gene and that those pairs are more often exclusions than co-mutations.

**Conclusions:** Based on this properties, we propose new drivers that warrant experimental validation. Our analysis is potentially relevant for cancer biology and classification, as well as for the rational selection of multitargeted therapeutic approaches.

## Background

Cancer is one of the most important health problems worldwide and, despite recent advances in diagnosis and therapy, cancer-associated mortality remains unacceptably high. Malignant tumors originate from genomic and epigenomic modifications which lead to loss of control of the cellular circuitry. Alteration of specific pathways enables tumors to bypass or activate a particular set of cellular processes, the so-called hallmarks of cancer [1], that confer tumor cells with adaptative advantages.

A particular biological pathway in cancer cells can be altered by somatic mutations or other changes in several genes. For example, in glioblastoma multiforme (GBM) the p53 pathway is downregulated in up to 87% of the tumors; but the genetic basis of this downregulation varies from patient to patient being the possible causes: somatic mutations or homozygous deletion of the protein p53 (TP53) or the cyclin dependent kinase inhibitor 2A (CDKN2A) and amplification of two genes codifying the double minute proteins (MDM2/MDM4). This and other examples have provided increased evidence that genetic alterations in cancer-related genes cluster within a limited set of essential biological pathways [2, 3, 4].

Tumor profiling projects have also unveiled mutually exclusive alterations across many patients, including driver mutations in specific genes [5]. For instance, TP53 mutations and MDM2 amplification in GBM very uncommonly occur together (few patients harbor both lesions). Additional examples include the mutual exclusivity in colorectal cancer between mutations in the adenomatous polyposis coli protein (APC) and catenin beta-1 (CTNNB1) genes (both involved in the beta-catenin signaling pathway) or mutations in BRAF and KRAS (genes of the RAS/RAF signaling pathway). In serous ovarian cancer, a mutual exclusivity between mutations of the breast cancer type susceptibility proteins BRCA and BRCA2 and the epigenetic silencing of BRCA1 has been observed while mutations in EGFR and KRAS are mutually exclusive in non-small lung cancer [2].

But cancer profiling has also discovered several cases of co-occurring alterations, suggesting that some changes in associated pathways may elicit complementary rather than redundant effects [6]. Examples include the PTEN (Phosphatidylinositol 3,4,5-trisphosphate 3-phosphatase and dual-specificity protein phosphatase) deletion concomitant with ERBB2 (Receptor tyrosine-protein kinase) amplification in breast cancer [7], MET (Hepatocyte growth factor receptor) activating mutations when VHL (Von Hippel-Lindau disease tumor suppressor) is deleted in renal carcinoma [8] and the CDKN2A suppression together with BRAF activating mutations in melanoma [9, 2].

Despite all this evidence, to the best of our knowledge, there are not systematic analysis in the literature of co-dependent mutations across all types of cancer. The only studies available, which used the TCGA dataset and the 2008 version of COSMIC, deal with whole genes and not specific mutations [10, 2, 11, 12, 13, 14], ignoring the fact that different mutations in the same gene can have widely different effects, i.e. G735S, G796S and E804G induce oncogenic activation of EGFR in prostate cancer while R841K has no functional relevance [15].

A better and wider understanding of co-dependencies between mutations is relevant in many aspects, such as tumour classification, diagnosis or treatment choice. At this respect, co-dependency relationships between genetic alterations evidence mutational epistasis [16] and highlight the need for multitargeted therapy. Several new generation antitumor drugs target proteins carrying specific driver mutations, i.e. sorafenib is active against renal and hepatic cell carcinomas harboring the BRAF.V600E mutation [9]; imatinib against gastrointestinal stromal tumors with mutations V560G, K642E, N822H or N822K in KIT or mutation V561D in PDGFRA [17]; gefitinib, erlotinib and afatinib against non-small cell lung cancers with exon 19 deletions or L858R in EGFR [18]; and dabrafenib or vemurafenib against BRAF.V600E in melanoma [19]. However, these treatments focusing in a single alteration are almost invariably followed by relapse due to selection of resistant cells [20]. Multitargeted approaches against co-occurring, biologically relevant mutations have the potential to delay the onset of resistance, and a better understanding of co-occurring oncogenic alterations could be of help in this setting. It must be remembered that some of these approaches are already in clinical use, such as the combination of BRAF and MEK inhibitors in metastatic melanoma [21] and many others are currently being tested in clinical trials [22].

Here we present the first systematic analysis of COSMIC somatic mutations [23] aimed to uncover cancer specific patterns of mutation associations, demonstrating that such an analysis is feasible and renders valuable information.

## Results

Data was downloaded from COSMICv75 and filtered to obtain a dataset of recurrent non synonymous mutated protein positions in common cancer types. A cancer type was defined as the unique combination of tissue, histology and sub-histology (e.g: lung /carcinoma /adenocarcinoma). Mutations were grouped by position in loci, e.g. BRAF.V600E, BRAF. V600D, BRAF.V600K and BRAF.V600R were grouped as BRAF.V600. Positions with driver mutations were identified according to Kin-driver database [24] in the case of protein kinases and to the literature in the case of NRAS, KRAS, HRAS [25, 26], PI3KCA [27] and TP53 proteins [28]. The dataset used for the analysis was composed of 1,098,411 samples from 687 cancer types with 365,096 mutated loci (289 of them are known driver loci) in 1,329 genes (see methods).

In order to avoid multi testing correction, previous approaches had focused on the identification of clusters of mutated genes through exploration of cellular pathways and statistical testing of significance [10, 2, 11, 12, 13, 14]. In contrast, we have used a simple pipeline combining multi testing with conditional and expected mutational probabilities to define pairs of co-dependent loci in the different types of cancer. Those pairs were subsequently merged in a single network where general traits about cancer co-mutation and exclusion could be observed.

### Counts of co-sequenced loci reveal pairs of significantly related mutations

As our first interest resided in related mutations, we tested all the pairs of co-sequenced loci where each member of the pair was mutated more than 10 times. This analysis was made in each cancer type, since admixing could hide signals characteristic of a particular malignancy. The arbitrary threshold of 10 mutated samples attempted to filter out uninteresting mutations that might co-occur randomly and to make the analysis more comprehensible. Thus, we obtained a starting dataset containing 262375 pairs of co-sequenced loci from a total of 135 cancer types.

Each pair of loci was tested for co-dependency using the exact Fisher test, which compares the number of samples with the two loci mutated with those with one or none. We found these numbers to be highly unbalanced due to the differential sequencing of loci -with some of them extensively and others only rarely sequenced-and the large proportion of samples with no mutations (supplementary figure 11). In consequence, false positives were possible; in particular, poorly co-sequenced pairs could be falsely detected as co-dependent owing to contingency tables with low values. To avoid this problem, we enhanced previous algorithms by combining the co-dependency tests with a comparison between observed and expected co-mutation probabilities (see methods “Filtering of probable false positives”). Thus, we ended up with 30,679 pairs of dependent loci in particular cancer types whose relation is not an artifact due to low mutational frequencies.

### Conditional probability discriminates the type of dependency between pairs of related loci

If the probability of mutation of a locus increased when another mutation in a different locus is present, we considered that those loci co-mutated. That is, if the a posteriori mutation probability of a locus surpassed its a priori mutation probability, a co-mutation was assigned in a particular cancer type; if the opposite happened, mutations were considered to be mutually exclusive. The difference between a priori and a posteriori probabilities was usually low; in consequence, we defined the confidence interval of the a priori probabilities for each mutation and subsequently classified the mutations according to the rules depicted in figure 1.

**Figure 1:**
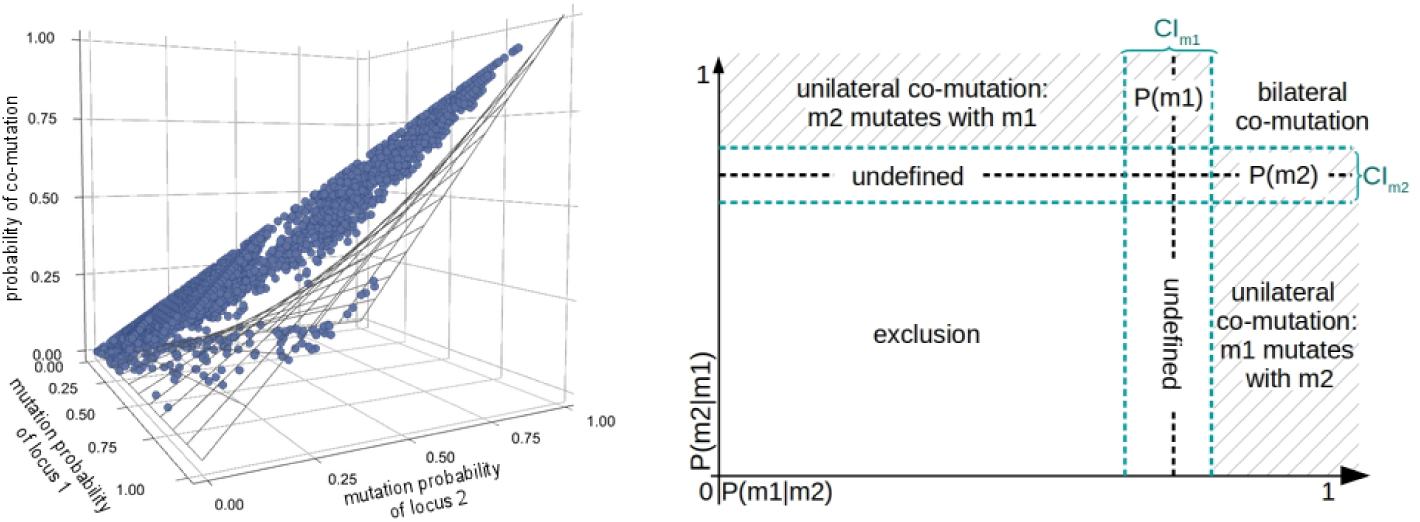
Treatment of dependent pairs. Left. Criteria used to filter possible false positives. Observed values are represented with blue balls, while the grey mesh follows the expected values assuming independence between loci. Pairs with an observed probability of co-mutation within the confidence interval of the expected one were discarded. Right. Rules used to classify the co-dependent pairs. The space of mutational probability of each pair is divided as shown by the relation between a priori P(*m*_*i*_) and a posteriori probabilities P(*m*_*i*_*|m*_*j*_).

If P(*m*_*i*_*|m*_*j*_) *< CI*(*m*_*i*_) AND P(*m*_*j*_*|m*_*i*_) *< CI*(*m*_*i*_) → mutual exclusion

If P(*m*_*i*_*|m*_*j*_) *> CI*(*m*_*i*_) AND P(*m*_*j*_*|m*_*i*_) *> CI*(*m*_*j*_) *→* bidirectional co-mutation

Where P(*m*_*i*_*|m*_*j*_) is the conditional probability of *m*_*i*_ given *m*_*j*_ and *CI*(*m*_*i*_) is the 95% confidence interval of the a priori probability of mutation of *m*_*i*_.

If the mutation probability of a locus augmented given a second mutation but the reciprocal didn’t happen, the co-mutation would have been considered unidirectional. However, in our analysis we didn’t encounter a single case of this class of dependency

### Network of loci with co-dependent mutations

As a result of our analysis, we found 189 pair of exclusions and 30,490 of bidirectional co-mutations (in total: 30,679 pairs of co-dependent mutations) involving 568 loci across 67 cancer types and 22 tissues of origin. The involvement of only 568 loci in the 30,679 pairs highlights the fact that a locus can mutate repeatedly in different types of cancer.

As 30,272 of the 30,679 pairs (98.67%) were found in upper aerodigestive tract/carcinoma/squamous cell carcinoma (supplementary figure 2, also interactive in http://sdmn.leloir.org.ar/, bottom link), they were studied separately (see below), and our analysis focused on the rest of the cancer types. Thus, we ended up with 407 pairs of related mutations, 218 co-mutations and 189 exclusions, involving a total of 260 loci from 94 genes in 66 cancer types. The pairs are shown in the network depicted in figure 2 and can be interactively explored in the following link: http://sdmn.leloir.org.ar/. The user can look for a locus (i.e: KRAS.G12) in the search box or see the dependencies it has by clicking the node. Also, by applying appropriate filters, users can see the relationships within a particular type of tumor or gene, display only the co-mutations, the exclusions, the genes listed in the Cancer Gene Census [29] and the driver mutations. In addition, Figure 2 integrates the mutational probability of each position (size of the node), the tissues where the pair occurs (edge color) and whether a protein pertains to the Census (black outline at the nodes) and if the locus is known to have driver mutations or not (red outline at the nodes). It is worth noting that we have found dependent mutations within 94 genes from a starting dataset of 1,329 genes, thus meaning that the remaining 1,235 genes have mutations that do not significantly associate with others.

**Figure 2:**
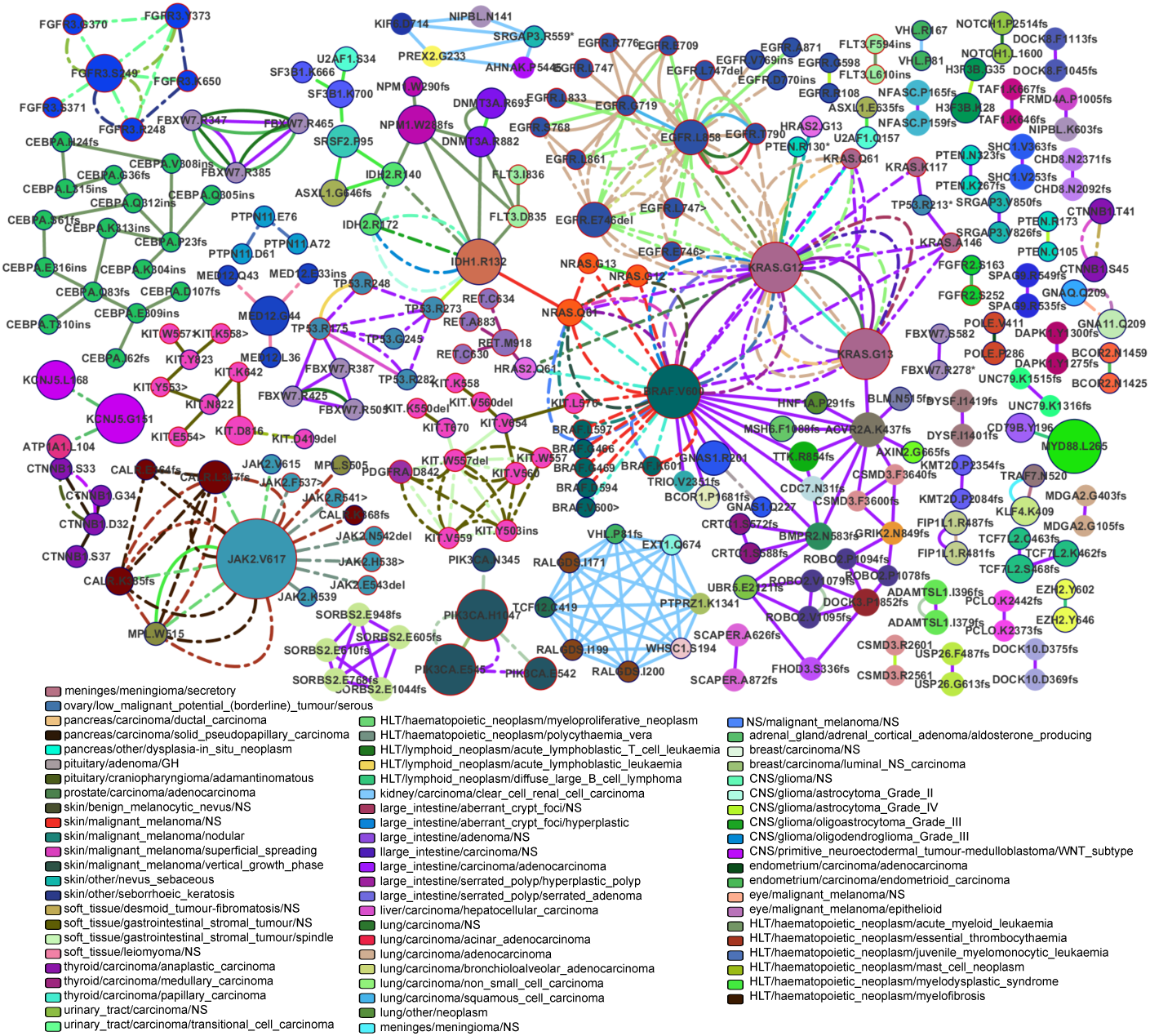
Network of significantly related loci. Also available for interactive view in: http://sdmn.leloir.org.ar/. Nodes are loci and edges are cancer types. The size of a node represents its mutational frequency. Nodes of the same colour pertain to the same protein. A border means that the protein is reported in the cancer Census. If the border is red, the locus has driver mutations. Two nodes are linked if they are significantly related in the cancer type of the corresponding color. Dotted lines indicate exclusions while continuous lines represent co-occurrences.

### Distribution of repeated pairs of mutations across cancer types and prevalence of proteins in some cancer types

The network depicted in figure 2 shows how some pairs of associated loci appear in different cancer types (nodes connected by more than one edge) and also how pairs of loci involving particular genes are prevalent in some tumors.

Regarding pairs repeated across different types of cancer, we found a total of 52, involving 58 loci from 24 proteins, all of them reported in the Cancer Gene Census. Additionally, of those 58 loci, 31 (57%) are known to harbor driver mutations. About two thirds (34) of the 52 pairs were found in different tumors from the same tissue of origin while 18 appeared in different tissues of origin. Although most of the 52 repeated pairs behaved equally across cancer types, we found a few examples where the type of association diverged; namely 6 pairs, which were mutually exclusive in most cancer types but occasionally co-occurred (see supplementary table 2).

We also found an abundance of specific genes in a particular type of tumor like CCAAT/enhancer-binding protein alpha (CEBPA) loci in 16 out of the 27 pairs found in acute myeloid leukaemia; KIT (Mast/stem cell growth factor receptor) loci as a partner in 19 out of 20 significantly related pairs of soft gastrointestinal stromal tumor (GIST); and 54 of 60 pairs in lung carcinomas involving loci from the epidermal growth factor receptor (EGFR). In fact, only large intestine/carcinoma/adenocarcinoma, the second cancer after upper aerodigestive tract/carcinoma/squamous cell carcinoma in number of pairs, escaped from this predominance of loci from specific proteins in the pairs in a particular type of cancer (figure 2 and supplementary figure 3a).

**Figure 3:**
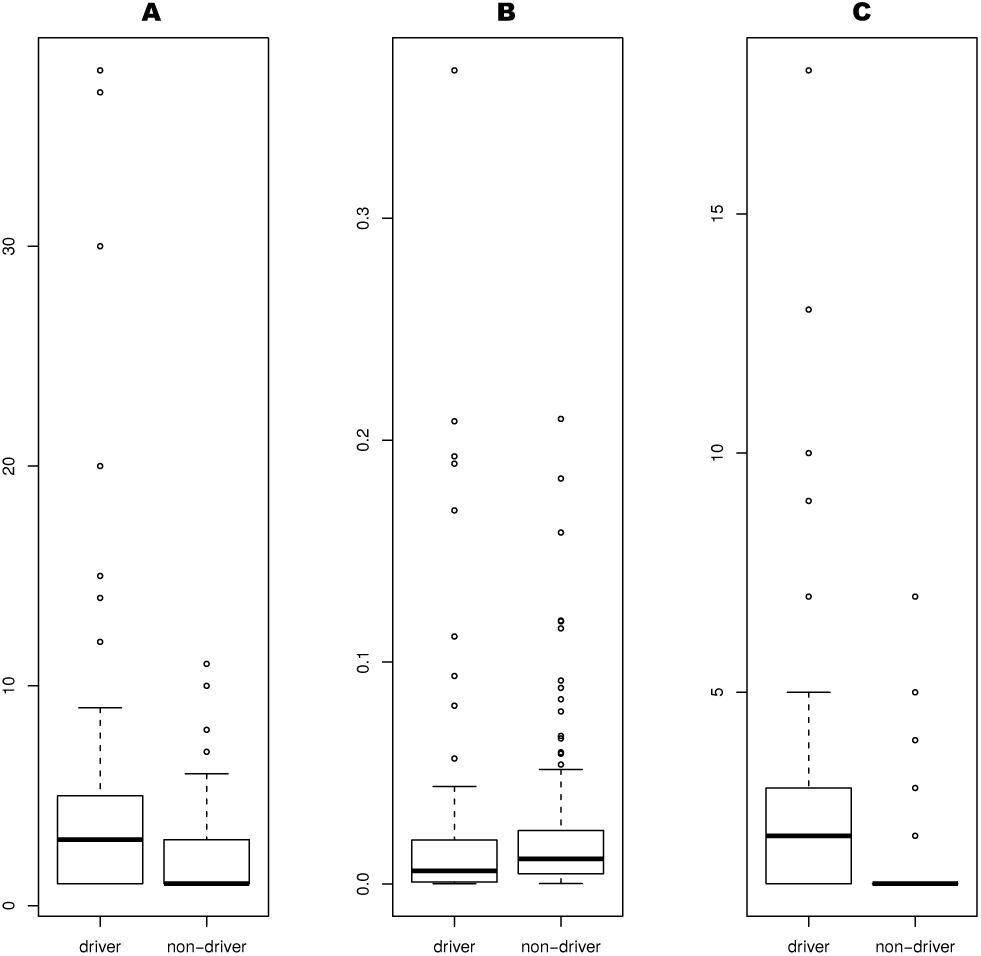
Difference between driver and non-driver loci. A. Connectivity. B. Mutational Frequency. C. Number of cancer types.

In our analysis we also found the presence of cliques, that is, groups of several interconnected loci, in the network of some tumors (all can be explored in interactive figure 2). Examples include the clique of exclusions between KIT.W557del, KIT.Y503ins, KIT.V559, KIT.V560, KIT.W557 and PDGFRA.D842 in soft tissue/gastrointestinal stromal tumour/NS and the clique formed by FGFR3.S249, FGFR3.Y373, FGFR3.G370 and FGFR3.R248 in urinary tract/carcinoma/transitional cell carcinoma. Other Interesting cliques were a group of 8 co-mutating loci involving 6 different proteins in kidney (kidney/carcinoma/clear cell renal cell carcinoma) and the 5 frameshifts co-occurring in SORBS2 in large intestine (see supplementary figure 3).

### Driver loci can be distinguished from non-driver loci based on their associations

Looking for specific properties of the driver loci (encircled with red in figure 2), we noticed a large quantity of edges of different tumors arising from common driver nodes like BRAF.V600 and KRAS.G12 (edges color stand for cancer types), suggesting that driver loci interact in a higher diversity of cancers than non-drivers. When we checked this analytically, we found that the two groups of nodes have a significantly distinct degree, mutational frequency and number of cancers in which they are present, as can be seen in figure 3. Driver loci have more edges (Mann Whitney U Test p-value = 1.583e-05) and are present in more cancer types (Mann Whitney U Test p-value = 8.86e-12) than non-drivers, but show lower mutational frequencies (Mann Whitney U Test p-value = 0.006581).

We were expecting that the loci more frequently mutated would be more connected in more types of cancer. But, while connectivity and number of cancers were highly correlated (Pearson coefficient: 0.8408, p-value = 2.2e-16), connectivity and mutational frequency and number of tumors affected and mutational frequency were only partially correlated (Pearson coefficient: 0.5205, p-value = 2.2e-16 and 0.4733, p-value = 6.3e-16, respectively).

### Loci with driver mutations tend to exclude while non-drivers tend to co-mutate

Next, we divided the pairs of related loci in those with (i) no driver, (ii) one driver and (iii) two driver loci. This classification was found to be associated with the type of relationship between the loci in the pair (X-squared p-value *<*2.2e-16). As shown in figure 4, the most common association between pairs of two driver loci is mutual exclusion, 80.5% (128/159). In contrast; 80.8% (151/188) of the no driver pairs were found to be co-mutations. It is worth noting that some well-known exclusions between driver loci appeared clearly in our analysis, such as the mutual exclusion of KRAS, BRAF and NRAS mutations in colorectal cancer [30](figure 2, zoomed in supplementary figure 3c).

**Figure 4:**
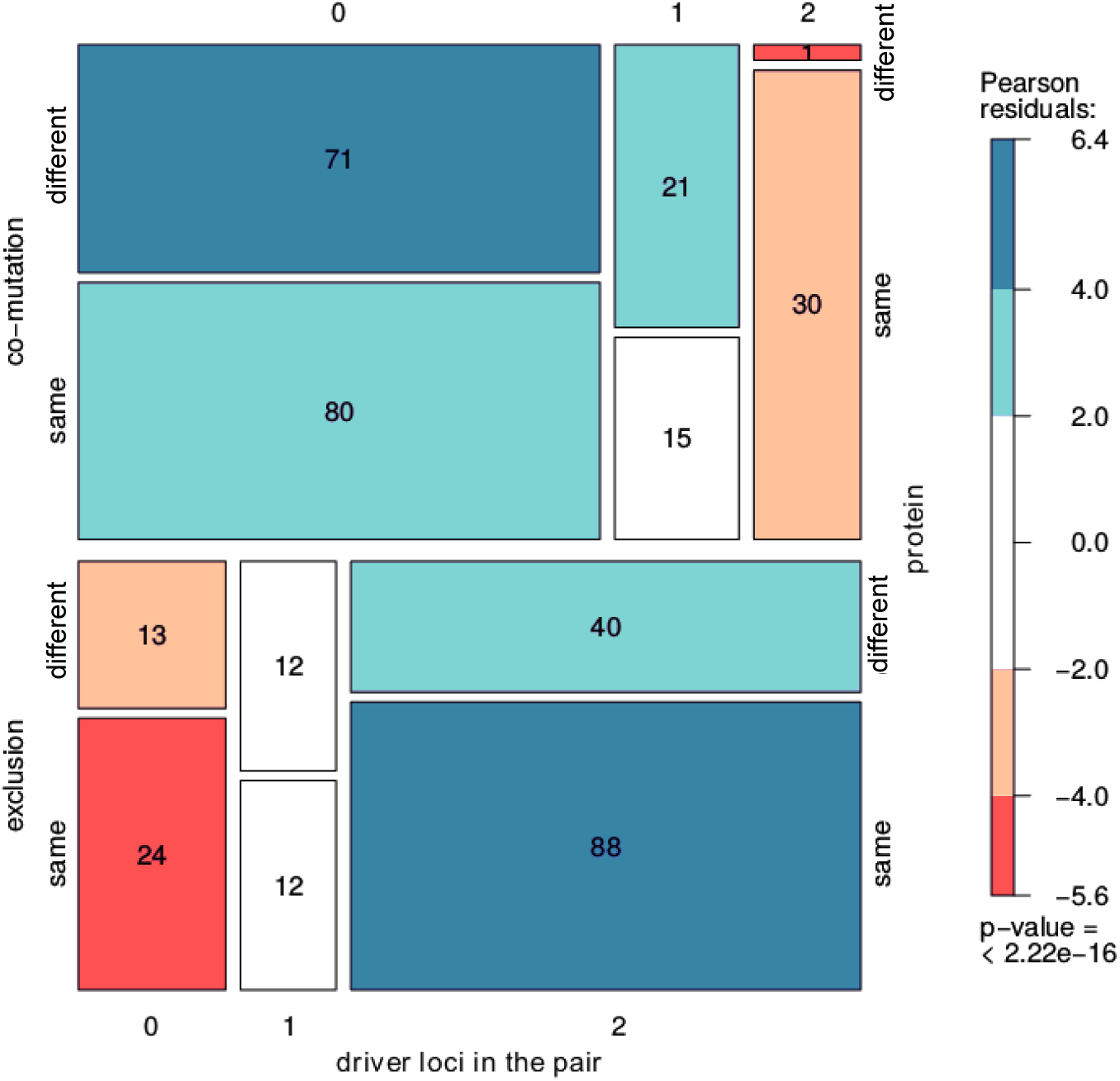
**Classification of pairs by the origin of the loci (different or same protein), the type of relation (co-mutation or exclusion) and the number of driver loci involved (0,1 or 2)**. Color indicates deviance from the expected under a model of complete independence. For example, pairs formed by 2 driver loci behave oppositely to pairs with no driver loci, with 128 exclusions versus 31 co-mutations. Also, they tend to involve same protein loci, with 88:40 pairs within the exclusions and 30:1 within co-mutations. The color of the box representing the co-mutation of driver loci from distinct proteins, shows that the value of 1 is between 4 and5.6 residuals under the expected value, so it is individually significant.

The only 20 co-mutations where both members are drivers comprises 25 loci from 7 proteins in 12 cancer types. Only one pair of associated loci are of two different genes, KRAS.G12 + BRAF.V600 in thyroid/carcinoma/anaplastic carcinoma. There are 6 pairs that appear in more than one cancer type, 5 of them involving EGFR in different types of lung carcinoma, with the co-mutation between EGFR.L858 and EGFR.T790 being the most common (present in 5 cancers).

### Driver mutation pairs tend to occur in the same gene

When we considered all pairs, we found loci of a single gene are associated almost as frequently as loci from two different genes (Mantel-Haenszel X-squared p-value =0.6578). Next, we applied the classification presented in Figure 4 and we found that pairs with no drivers and pairs with one driver do not show a preference to occur in the same or different genes. In contrast, pairs with 2 drivers behaved differently, with a majority of them associating loci from the same gene. Namely, 96.77% of driver co-mutations and 68.75% driver exclusions were found to involve loci in the same gene (Fisher Test p-value = 0.0009532; X-squared p-value = 0.002964).

### The squamous cell carcinoma of upper aerodigestive tract mutates in an uncommon way

Upper aerodigestive tract/carcinoma/squamous cell carcinoma (SCC) is the name used to denote a variety of cancers, with divergent etiologies, originated in the epithelium of head or neck [31]. It is by far the least sequenced malignancy in the COSMIC database, with 2048 samples versus the 32,392 of large intestine/carcinoma/adenocarcinoma (the most sequenced one). SCC exhibit a high mutational frequency that could explain the extremely elevated number of significant dependent pairs encountered (Supplementary figure 2, and http://sdmn.leloir.org.ar -bottom link-). The 30,272 pairs found in SCC correspond to co-mutations associating 307 loci of 45 proteins. It is compelling that there are no exclusion pairs and no previously described driver loci involved. Only 8 of the 45 proteins (PDE4DIP, NCOR1, HLA-A, NOTCH1, NOTCH2, BCOR1, KMT2C and SETBP1) are reported in the Census, forming 411 pairs with 65 loci

## Discussion

Although the advent of Next Generation Sequencing technologies has allowed the mutational profiling of thousands of tumor samples, no systematic studies have been published of co-occurring and mutually exclusive mutations. Here, we report a cancer-type specific bioinformatics analysis of the thousands of somatic mutations described in COSMIC, which were grouped in loci, aimed to discover co-occurrence and exclusion pairs. In our analysis, we found some well known pairs that validate our approach, such as the mutual exclusion of KRAS.G12/G13 with EGFR loci in lung cancer or KRAS.G12/13 with BRAF.V600 in several neoplasias. Another example is the co-ocurrence of EGFR.L858 + EGFR.T790 in lung adenocarcinoma. Somatic mutations in L858 confer sensitivity to tyrosine kinase inhibitors targeting EGFR (EGFR TKIs). However, patients ultimately relapse and one of the most common mechanisms of resistance to TKIs is the emergence of the p.T790M mutation. This observation constitutes an example of how a systematic analysis of co-ocurrences in tumor rebiopsies after progression to targeted therapies can help to find loci associated with resistance.

However, we also encountered controversial co-ocurrences like KRAS.G12 + KRAS.G13 in anaplastic thyroid carcinoma, prostate adenocarcinoma and papillary thyroid carcinoma or BRAF.V600 + KRAS.G12, again in anaplastic thyroid carcinoma. These mutations are generally regarded as mutually exclusive in most malignancies [32, 33]. To track this discrepancy, we reviewed the articles reporting co-mutations of this two loci. Garcia-Rostan et al, described co-mutations KRAS.G12 + KRAS.G13 in poorly differentiated (papillary) and undifferentiated (anaplastic) thyroid carcinomas, but did not discuss them further [34]. The same co-mutation but in prostate adenocarcinoma was reported in two articles [35, 32]; with Silan et al remarking the high Gleason Score and PSA (prostate specific antigen) levels on combinedly mutated patients, both being indicators of an aggressive tumor. Meanwhile, Costa et al identified a strong link between clinical parameters indicative of unfavourable prognosis and BRAF.V600E associated with other genetic events, such as the co-mutation BRAF.V600 + KRAS.G12 [36]. Another unexpected pair of associated loci was the co-mutation EGFR.L858 + EGFR.G719 in lung squamous cell carcinoma (LSCC), since the frequency of EGFR mutations in LSCC is low [37] and, as a consequence, the L858+G719 co-mutation is very uncommon, appearing only in approximately 1/2000 patients (frequency 0.0007520682).

Another issue that should be considered when trying to explain unexpected co-mutations is intratumoral heterogeneity. Subclonal populations within the same tumor [38] can explain the presence of mutually exclusive mutations, as sequencing admixes the genomes of different cells. Zou et al even report a BRAF mutation in a whole thyroid tumor, and RAS mutation only in some sections [39]. This examples of tumor heterogeneity reflect the need for serial examination of tumors during the course of therapy, and of different areas within a single tumor [40]. Genetic heterogeneity has also been linked to a variable clinical response to treatment, with primary tumors and metastatic regions responding differently to the same drug [40]. Whichever is the explanation for the unexpected co-mutations, they are probably capturing a general feature of the corresponding tumors that warrants testing and, if found, should be considered during treatment. In tumors with BRAF.V600 and KRAS.G12, either via a true co-mutation or due to tumor heterogeneity, both loci should be targeted to avoid the selection of subclonal resistance mutations through treatment [38].

During our search for mutational patterns, we made a number of general observations regarding pairs of associated loci. First, we discovered that loci within a single gene (or occasionally two genes) are present in a particular type of cancer, indicating a relevant role for the corresponding protein(s) in that specific malignancy. Some examples were CEBPA in acute myeloid leukemia, EGFR and KRAS in lung cancer and KIT in GIST. According to the literature activating KIT mutations are present in a majority of GISTs from soft tissue [41], while the oncogenes EGFR and KRAS in lung cancer, are not always mutated but serve for routinary diagnoses [42, 43]. Large intestine adenocarcinoma was the only relevant tumor that did not show an association with a particular gene in the co-mutational and exclusion patterns. This finding was not unexpected, since it has been demonstrated that several genes can play key roles in the development of this malignancy [44, 45].

We also observed that driver mutations present three common properties, namely (i) driver loci tend to mutate in a mutually exclusive fashion, (ii) driver loci pairs are repeatedly present in several cancer types and (iii) driver loci pairs frequently occur within the same gene. These three properties can facilitate the discovery of new drivers and, as a proof of concept of this idea, we searched loci fulfilling at least one property in our network of associations, finding a total of 172 possible new driver loci in genes of the Cancer Census (supplementary table 3). Of them, 15 were located in 4 protein kinases, which we further analyzed via structural alignment using the Kin-driver database. We found that 8 loci were in positions known to be drivers in other kinases, and 9 mapped to hyper mutated segments (-HS-) where driver mutations have been shown to cluster [46] (Table 1). Furthermore, one of these loci, KIT.D419del, has indeed been described as a driver [47] although it was not considered as such in our analysis; and there is experimental evidence suggesting a driver role for 7 additional loci [48, 49, 50, 51, 52].

**Table 1:**
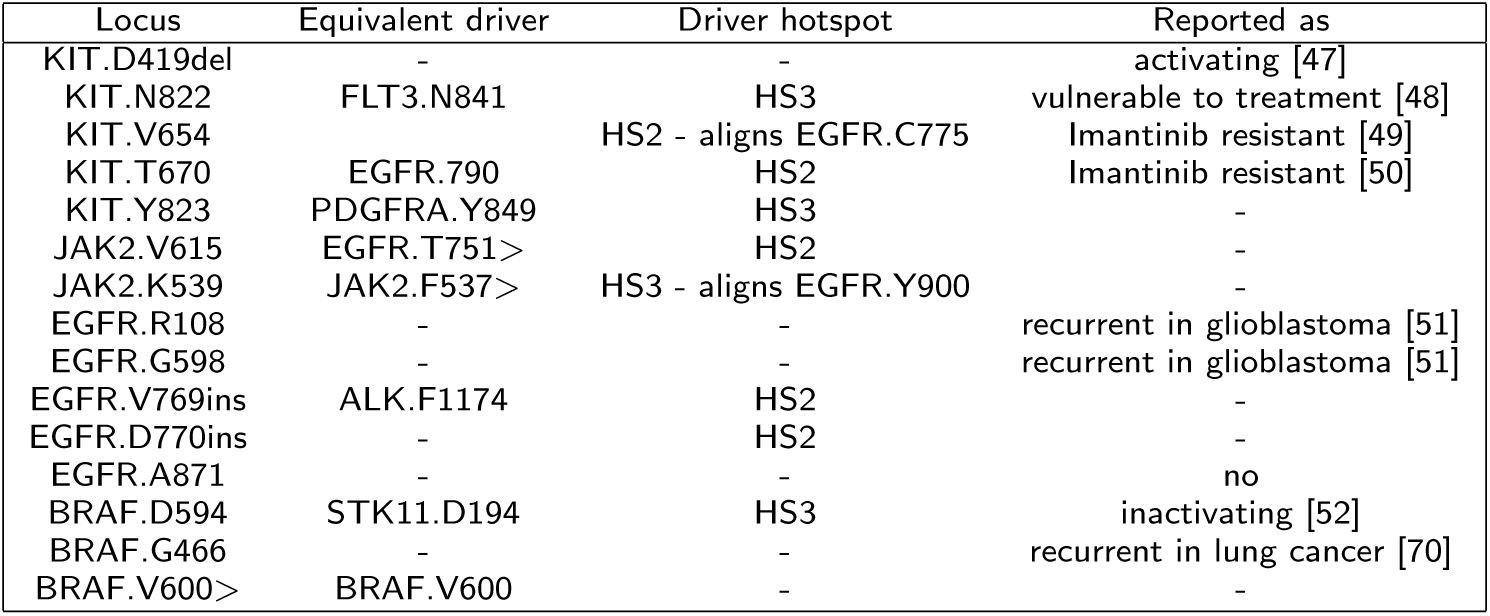
Kinases loci suggested as driver by the analyses

In addition to protein kinases, some loci in other types of genes exhibited the three properties mentioned above so we consider them as candidate drivers. Examples include R282 in the TP53 gene or K385fs and L367fs in the calreticulin gene (CALR). Mutations in the TP53.R282 locus are relatively common and they exclude with the drivers TP53.R175 and TP53.R273 in two cancer types according to our results. According to literature mutation of R282W is actually a driver, it has been described to shift the DNA binding domain of the p53 protein to a dysfunctional structure [53] and cause an earlier onset of familial cancer and a shorter overall survival in cancer patients [54]. Regarding CALR. L367fs and CALR.K385fs, they exclude with other frameshifts in the same gene, and also with thrombopoietin receptor (MPL) V515 and the know driver JAK2.V617 in three haematopoietic neoplasms. This pattern is not a new finding; the triad of exclusion amidst JAK2, MPL and CALR has been previously reported [55]. Frameshifts in exon 9 of CALR gene, such as CALR.L367fs and CALR.K385fs, are common in some myeloproliferative disorders and change the C terminal charge of the protein, altering its subcellular localization, stability and function [56, 57].

In addition to pairs, we found cliques of significantly related loci (listed in supplementary Table 4). The co-mutation cliques, where mutations in each loci are more probable when some of the other loci are mutated, can represent combinations conferring synergistic adaptative advantages to the tumor cells. An example is the clique formed by IDH1.R132, NPM1.W288fs, DNMT3A.R882 and FLT3.D835 in acute myeloid leukaemia that emerged in our analysis. Mutations in IDH1 and DNMT3A, and in DNMT3A and FLT3, have been described to be simultaneously present in a significant number of myeloid leukaemia patients, and concomitant mutations in the triad NPM1/DNMT3A/FLT3 have been associated with a worse overall survival in this malignancy [58, 59]. These experimental findings support our hypothesis of a synergistic advantage of co-mutation cliques. In contrast, exclusion cliques, probably reveal loci that alter the same biological pathway. A mutation in one of them is enough to acquire the corresponding hallmark of cancer and, therefore, tends to exclude the others [6].

The pairs and cliques of associated loci that emerged in our analysis, particularly those involving drivers, might prove useful not only in cancer biology studies, but also for the selection of therapies. Cancer treatment faces several challenges, such as the selection of appropriate markers for targeted and non-targeted agents and the relapse to a more aggressive disease after an initially successful treatment. Since most of the new antitumor drugs specifically target mutated or genetically altered proteins, co-mutations can suggest combined treatments that can prove more effective than single agent approaches. In contrast, exclusions might indicate that certain combinations of drugs are unlikely to be useful in a meaningful percentage of patients. For instance, we found that the IDH1.R132 and the TP53.R273 loci co-mutated in gliomas. Mutations in IDH1.R132 gene are very frequent in this malignancy and inhibitors of mutant IDH are currently in trials to prove their clinical utility as single agent or in combination strategies targeting additional oncogenic pathways [60]. Anti-mutant TP53 drugs are also being tested in clinical trials [61], and our results indicate that they might be an appropriate partner for IDH inhibitors in a significant number of gliomas.

The absence of links is also of interest. One example are mutations in PTEN that lead to the loss of protein expression and have been related to resistance to many targeted therapies, such as EGFR inhibitors in EGFR-mutant lung cancer [62], anti-EGFR antibodies in non mutated KRAS/NRAS colorectal cancer [63] or BRAF inhibitors in BRAF-mutant melanoma [64]. In our analysis, PTEN mutations are not significantly dependent to EGFR, KRAS, NRAS or BRAF mutations in different cancer types with the only exception of the co-mutation with KRAS.G12 in endometrium/carcinoma shown in the figure 2. In consequence, it can be estimated that, if the above-mentioned therapies are tested in patients of all the other tumor types, the percentage of cases not responding due to loss of PTEN will be equal to the overall frequency of PTEN mutations in that particular malignancy. Data of this kind can be of great interest when trying to find new applications for targeted agents.

One important limitation of our study derives from the fact that a vast majority of the tumors compiled in the COSMIC database have only sequenced a limited number of genes, while whole exome or whole genome sequenced tumors are scarce. In consequence, we are likely missing a significant number of associations between loci simply because they have been rarely sequenced. This limitation also explains a counterintuitive correlation found during our analysis, where the more mutated loci are not the more connected ones (Figure 3). For example, the G151 mutation of the potassium channel KCNJ5 was almost as frequently mutated in the samples of the COSMIC database as KRAS.G12, with values of 0.1827 and 0.1893, respectively; but KCNJ5.G151 was significantly less connected. The explanation lies in the fact that, while the ubiquitous driver (KRAS.G12) has been sequenced on 164511 samples from all types of tumor, the KCNJ5 status has only been reported in 2671 samples, most of them (81.2%) from adrenal gland/adrenal cortical adenoma/aldosterone producing, a cancer type where KCNJ5 mutations are prevalent [65]. To overcome this limitation, we plan to periodically update our analysis and we are confident that we will find new associations of loci relevant for both cancer biology and treatment.

The three-properties approach, might not be adequate to find possible drivers loci in upper aerodigestive tract squamous cell carcinoma. Driver mutations for this cancer may have exceptional characteristics or, more likely, drivers in this malignancy are genomic aberrations different from mutations. Experimental evidence seems to support this explanation, since non mutation drivers such as EGFR overexpression and amplification have been described to be frequent in this malignancy [66].

## Conclusions

In summary, we can propose driver mutations based solely on the network of significantly related pairs of mutations. At now, driver predictions rely on interaction and functional networks focusing on complete genes [67, 68, 69], then, our approach have the advantage of pinpoint specific mutated positions, which enlightens the functional role they may be playing. All this prove the relevance of cumulative repositories like COSMIC and cBio, that aggregate enough data sets to search for significant patterns.

## Methods

### Dataset

Complete COSMICv75 was downloaded. Data included 1,178,444 samples from 47 tissues with 193 histologies and 716 sub-histologies; there were 2,812,088 mutations from 2,128,846 sequenced positions. To identify unique mutations, ENSEMBL transcript ID from COSMIC entries were mapped to UNIPROT protein ID and concatenated with the mutations (e.g P15056.V600E). Mutations were grouped by position following the type of alteration, so we could distinguish substitutions, deletions, insertions, complex substitutions and frameshifts.

Mutations of type Nonstop extension, Substitution coding silent and Whole gene deletion were filtered. This way the dataset decreased to 1,107,460 samples from 1,298 cancer types with 1,615,508 mutated positions from 19,297 proteins. Among these, 291 positions with driver mutations in kinases, Ras proteins, TP53 or PIK3CA were found. Cancer type was defined as the unique combination of tissue, histology and sub-histology (e.g: lung/carcinoma/adenocarcinoma).

In order to roughly discard unimportant (to cancer evolution) mutations, positions sequenced in less than 1,000 samples were filtered. The threshold is set empirically to a limit passed by most of the driver loci (289 from 291) but only by 22.60% of the loci without known driver mutations. Cancer types without mutations or sequenced less than 10 times, were also discarded. Then, the dataset for the analysis is composed by 1,098,411 samples from 687 cancer types 365,096 mutated positions (289 of them are driver mutations) of 1,329 proteins.

### Co-dependant pairs of loci

To identify significantly related mutations, contingency tables were constructed for every pair of co-sequenced loci in each cancer type. To avoid very infrequent mutations, only the pairs with more than 10 mutated samples (out of 1000 or more sequenced samples) for each position were retained, giving a starting dataset of 262,375 tables (pair of mutations to be evaluated) of 135 cancer types. Each table was checked for dependency between loci with Fisher test. P-values were FDR corrected. A total of 44,250 pairs of loci were found to be significantly associated in 82 cancer types.

### Filtering of probable false positives

As we feared that low mutational probabilities allowed unrelated pairs to pass the corrected Fisher test, we compared the expected and the observed probabilities for each pair. Specifically, scarcely mutated loci could seem to be excluding each other when they are just infrequent, so their expected joint probability would be similar to the observed one.

To filter this kind of false positives, we estimated the 99% confidence interval of each observed joint mutational probability and located the expected joint mutational probability. If the expected fell within the confidence interval of the observed, the pair was discarded, which is equivalent to filter all the dots near to the diagonal
in the figure 1.

Confidence intervals were calculated using the binomial distribution. The distribution B(n, p) is defined for each pair with size n, as the total of samples with both loci sequenced and probability of success p, as the observed probability of co-mutation.

After this step a final dataset of 30,679 pairs of dependent loci whose relation is not due to their low mutational frequencies was obtained.

### Assignation of type of dependency

To distinguish pairs that co-mutate from pairs that exclude mutually, a priori and a posteriori mutation probabilities were compared. Considering only samples with loci A and B co-sequenced, if locus A has a higher probability of mutation when mutation B is present than in the whole set of samples, A co-mutates with B. If the contrary is true, mutations exclude themselves. But the a priori mutational probability of a locus can vary randomly, so we estimated its 95% confidence interval as previously described. This way, each table defines its own thresholds but a coherent set of simple logic rules are used over the whole dataset:

If P(*m*_*i*_*|m*_*j*_) *< CI*(*m*_*i*_) AND P(*m*_*j*_*|m*_*i*_) *< CI*(*m*_*i*_) → mutual exclusion

If P(*m*_*i*_*|m*_*j*_) *> CI*(*m*_*i*_) AND P(*m*_*j*_*|m*_*i*_) *> CI*(*m*_*j*_) →bidirectional co-mutation

Where P(*m*_*i*_*|m*_*j*_) is the conditional probability of *m*_*i*_ given *m*_*j*_ and CI(*m*_*i*_) is the 95% confidence interval of the unconditional probability of mutation of *m*_*i*_.

### Network of related loci

The 30,679 pairs were divided by the type of dependency in 189 pairs of exclusion and 30,490 of co-mutation. Upper aerodigestive tract/carcinoma/squamous cell carcinoma pairs were separated and two network representations were made in cytoscape. The one for the upper aerodigestive tract squamous cell carcinoma was ordered with the prefuse force directed layout based on the observed probability of co-mutation and enhanced manually to show all node labels. The other network, representing the rest of the pairs, also was enhanced manually but departing from an orthogonal layout.

### Properties of driver loci

Degree, mutational frequency and number of engaged cancer types was calculated for each loci in the network. The number of engaged cancer types was normalized by the maximum possible, 66 cancers. Distributions of the four variables (degree, mutational frequency, number of engaged cancers and normalized number of cancers) were compared between drivers and non-driver with Mann Whitney tests. The relation among the variables was tested with Pearson correlation.

Pairs were classified in a 3-level contingency table by their categorical variables: type of relation within the loci, origin of the linked loci, and number of driver positions involved. This was plotted in a mosaic and tested for dependency with Mantel-Haenszel test. Afterwards the segment of the table counting pairs formed by two drivers was tested with chi square.

Loci corresponding to proteins of the Cancer Genes Census and satisfying at least one of the observed properties of drivers were suggested as new possible drivers. The new possible drivers belonging to kinases were mapped to driver hotspot regions or to previously described driver loci. All calculus were made in R. The packages used are: igraph, Matrix, doParallel and foreach.

### Declarations

#### Ethics approval and consent to participate

Not applicable

#### Consent for publication

Not applicable

#### Availability of data and materials

The dataset supporting the conclusions of this article is included within the article and its additional files.

#### Author’s contributions

SO and CMB: had the idea, results generation, data analysis and discussion and manuscript writing. DJZ: statistics, EMP: web development, MAMV: assessor, discussions, writing the manuscript.

## Acknowledgements

We thank Dr. Octavio Arizmendi Echegaray for its help with the statistical design. SOM: has a fellowship from CONCYTEG, CMB and DJZ is a researcher of the National research council (CONICET), EMP has an “agencia” (MinCyT) fellowship. The work was supported by prestamo BID PICT2014-1787.

## Author details

^1^Fundación Instituto Leloir, Avda. Patricias Argentinas, C1405BWE Buenos Aires, Argentina. ^2^Laboratorio de Oncología/Pangaea Oncology, Hospital Universitario Quirón Dexeus, C. Sabino Arana 5, 08023 Barcelona, Spain.

---

**Supplementary figure 1:**
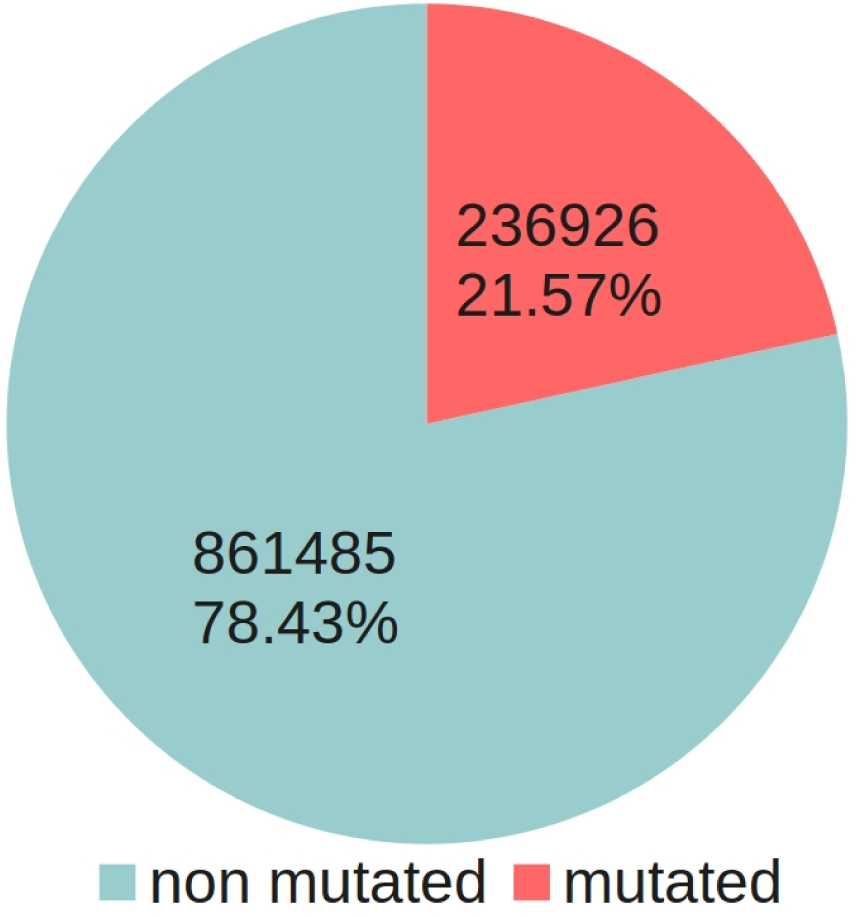
Type of samples in the dataset. There is almost three times more non mutated samples than mutated ones.

**Supplementary figure 2:**
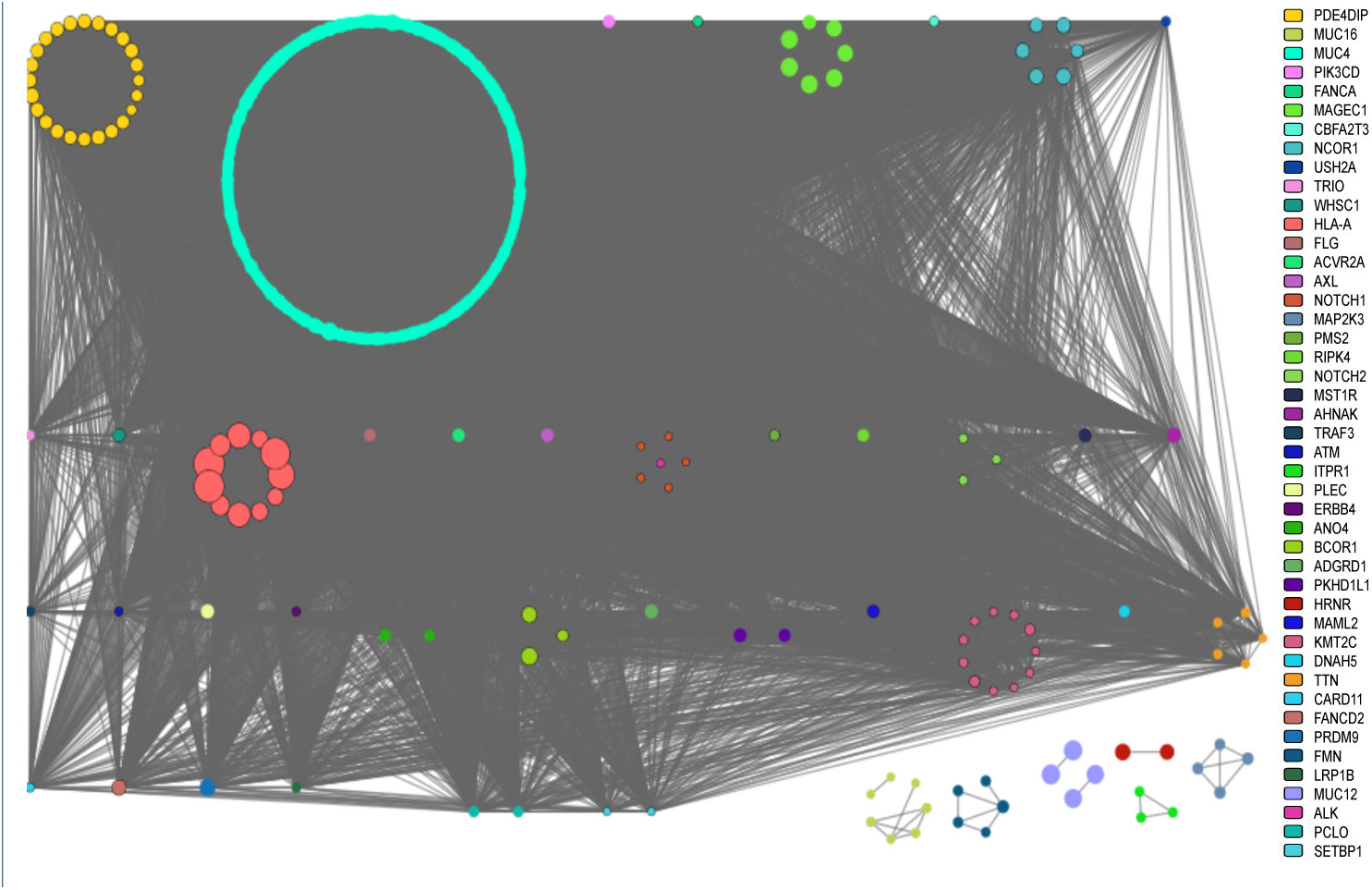
Dependent pairs of loci from upper aerodigestive tract/carcinoma/squamous cell carcinoma. Loci are represented by nodes, they are colored according to the protein of provenance and linked if they co-mutate. Node size reflects mutational frequency.

**Supplementary figure 3:**
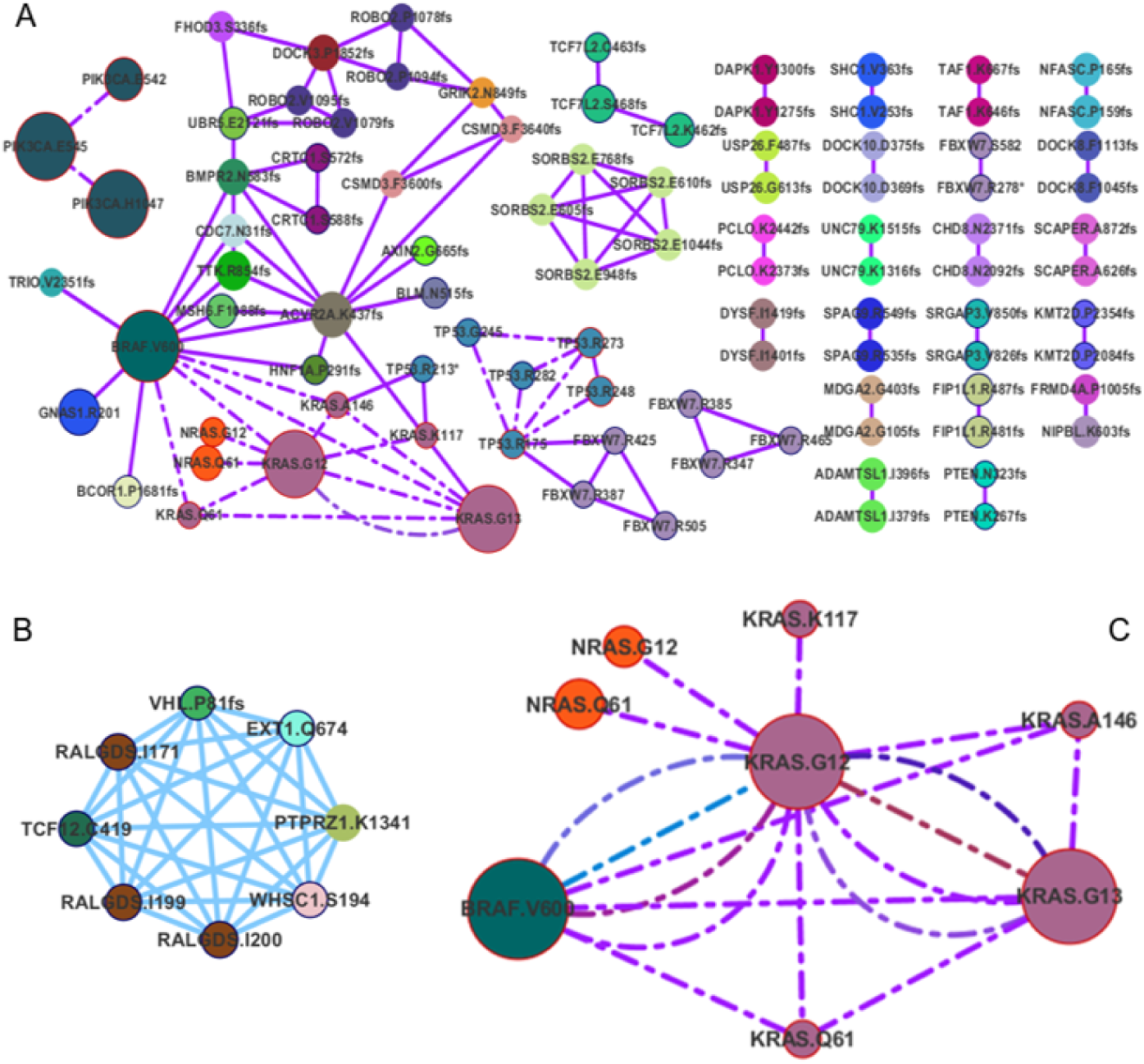
Extracts of the network of dependent loci. A. Dependencies in large intestine. Contrary to what happens in most cancer types, there is no protein predominantly represented. B. Clique of co-mutations in kidney, involving 6 different proteins. D. NRAS, KRAS and BRAF exclusions in large intestine.

## References

[1] Hanahan, D., Weinberg, R.A.: Hallmarks of cancer: The next generation. Cell 144(5), 646–674(2011). doi:10.1016/j.cell.2011.02.013

[2] Ciriello, G., Cerami, E., Sander, C., Schultz, N.: Mutual exclusivity analysis identifies oncogenic network modules. Genome Research 22(2), 398–406(2011). doi:10.1101/gr.125567.111

[3] Jones, S., Zhang, X., Parsons, D.W., Lin, J.C.-H., Leary, R.J., Angenendt, P., Mankoo, P., Carter, H., Kamiyama, H., Jimeno, A., et al.: Core signaling pathways in human pancreatic cancers revealed by global genomic analyses. Science 321(5897), 1801–1806(2008)

[4] Compagno, M., Lim, W.K., Grunn, A., Nandula, S.V., Brahmachary, M., Shen, Q., Bertoni, F., Ponzoni, M., Scandurra, M., Califano, A., Bhagat, G., Chadburn, A., Dalla-Favera, R., Pasqualucci, L.: Mutations of multiple genes cause deregulation of nf-kappab in diffuse large b-cell lymphoma. Nature 459, 717–721(2009). doi:10.1038/nature07968

[5] Kim, Y.-A., Madan, S., Przytycka, T.M.: Wesme: uncovering mutual exclusivity of cancer drivers and beyond. Bioinformatics, 242 (2016). doi:10.1093/bioinformatics/btw242

[6] Thomas, R.K., Baker, A.C., DeBiasi, R.M., Winckler, W., LaFramboise, T., Lin, W.M., Wang, M., Feng, W., Zander, T., MacConnaill, L.E., et al.: High-throughput oncogene mutation profiling in human cancer. Nat Genet 39(3), 347–351(2007). doi:10.1038/ng1975

[7] Teschendorff, A.E., Caldas, C.: The breast cancer somatic “muta-ome”: tackling the complexity. Breast Cancer Res 11(2), 301 (2009). doi:10.1186/bcr2236

[8] Bommi-Reddy, A., Almeciga, I., Sawyer, J., Geisen, C., Li, W., Harlow, E., Kaelin, W.G., Grueneberg, D.A.: Kinase requirements in human cells: Iii. altered kinase requirements in vhl-/- cancer cells detected in a pilot synthetic lethal screen. Proceedings of the National Academy of Sciences 105(43), 16484–16489(2008). doi:10.1073/pnas.0806574105

[9] Pratilas, C.A., Xing, F., Solit, D.B.: Targeting oncogenic braf in human cancer. Current Topics in Microbiology and Immunology, 83–98(2010)

[10] Babur, O., Gönen, M., Aksoy, B.A., Schultz, N., Ciriello, G., Sander, C., Demir, E.: Systematic identification of cancer driving signaling pathways based on mutual exclusivity of genomic alterations. Genome Biology 16(1) (2015). doi:10.1186/s13059-015-0612-6

[11] Kim, Y.-A., Cho, D.-Y., Dao, P., Przytycka, T.M.: Memcover: integrated analysis of mutual exclusivity and functional network reveals dysregulated pathways across multiple cancer types. Bioinformatics 31(12), 284–292(2015). doi:10.1093/bioinformatics/btv247

[12] Rubio-Perez, C., Tamborero, D., Schroeder, M.P., Antolín, A.A., Deu-Pons, J., Perez-Llamas, C., Mestres, J., Gonzalez-Perez, A., Lopez-Bigas, N.: In silico prescription of anticancer drugs to cohorts of 28 Tumor types reveals targeting opportunities. Cancer Cell 27(3), 382–396(2015). doi:10.1016/j.ccell.2015.02.007

[13] Cui, Q.: A network of cancer genes with co-occurring and anti-co-occurring mutations. PLoS ONE 5(10), 13180 (2010). doi:10.1371/journal.pone.0013180

[14] Yeang, C.-H., McCormick, F., Levine, A.: Combinatorial patterns of somatic gene mutations in cancer. The FASEB Journal 22(8), 2605–2622(2008). doi:10.1096/fj.08-108985

[15] Cai, C.Q., Peng, Y., Buckley, M.T., Wei, J., Chen, F., Liebes, L., Gerald, W.L., Pincus, M.R., Osman, I., Lee, P.: Epidermal growth factor receptor activation in prostate cancer by three novel missense mutations. Oncogene 27(22), 3201–3210(2008). doi:10.1038/sj.onc.1210983

[16] Nussinov, R., Tsai, C.-J.: “latent drivers” expand the cancer mutational landscape. Current Opinion in Structural Biology 32, 25–32(2015). doi:10.1016/j.sbi.2015.01.004

[17] Heinrich, M.C.: Kinase mutations and imatinib response in patients with metastatic gastrointestinal stromal tumor. Journal of Clinical Oncology 21(23), 4342–4349(2003). doi:10.1200/jco.2003.04.190

[18] Wang, X., Goldstein, D., Crowe, P., Yang, J.-L.: Next-generation egfr/her tyrosine kinase inhibitors for the treatment of patients with non-small-cell lung cancer harboring ¡em¿egfr¡/em¿ mutations: a review of the evidence. OncoTargets and Therapy Volume 9, 5461–5473(2016). doi:10.2147/ott.s94745

[19] Gençler, B., Gönül, M.: Cutaneous side effects of braf inhibitors in advanced melanoma: Review of the literature. Dermatology Research and Practice 2016, 1–6(2016). doi:10.1155/2016/5361569

[20] Tian, T., Olson, S., Whitacre, J.M., Harding, A.: The origins of cancer robustness and evolvability. Integr. Biol. 3(1), 17–30(2011). doi:10.1039/c0ib00046a

[21] Simeone, E., Grimaldi, A.M., Festino, L., Vanella, V., Palla, M., Ascierto, P.A.: Combination treatment of patients with braf-mutant melanoma: A new standard of care. BioDrugs : clinical immunotherapeutics, biopharmaceuticals and gene therapy 31, 51–61(2017). doi:10.1007/s40259-016-0208-z

[22] Bayat Mokhtari, R., Homayouni, T.S., Baluch, N., Morgatskaya, E., Kumar, S., Das, B., Yeger, H.: Combination therapy in combating cancer. Oncotarget 8, 38022–38043(2017). doi:10.18632/oncotarget.16723

[23] Forbes, S.A., Bindal, N., Bamford, S., Cole, C., Kok, C.Y., Beare, D., Jia, M., Shepherd, R., Leung, K., Menzies, A., et al.: Cosmic: mining complete cancer genomes in the catalogue of somatic mutations in cancer. Nucleic Acids Research 39(Database), 945–950(2010). doi:10.1093/nar/gkq929

[24] Simonetti, F.L., Tornador, C., Nabau-Moreto, N., Molina-Vila, M.A., Marino-Buslje, C.: Kin-driver: a database of driver mutations in protein kinases. Database 2014(0), 104–104(2014). doi:10.1093/database/bau104

[25] Rajagopalan, H., Bardelli, A., Lengauer, C., Kinzler, K.W., Vogelstein, B., Velculescu, V.E.: Tumorigenesis: Raf/ras oncogenes and mismatch-repair status. Nature 418(6901), 934–934(2002). doi:10.1038/418934a

[26] Moura, M.M., Cavaco, B.M., Leite, V.: Ras proto-oncogene in medullary thyroid carcinoma. Endocrine-Related Cancer 22(5), 235–252(2015). doi:10.1530/erc-15-0070

[27] Pao, W., Girard, N.: New driver mutations in non-small-cell lung cancer. The Lancet Oncology 12(2), 175–180(2011). doi:10.1016/s1470-2045(10)70087-5

[28] Sherr, C.J.: Principles of tumor suppression. Cell 116(2), 235–246(2004). doi:10.1016/s0092-8674(03)01075-4

[29] Futreal, P.A., Coin, L., Marshall, M., Down, T., Hubbard, T., Wooster, R., Rahman, N., Stratton, M.R.: A census of human cancer genes. Nat Rev Cancer 4(3), 177–183(2004). doi:10.1038/nrc1299

[30] De Roock, W., Claes, B., Bernasconi, D., De Schutter, J., Biesmans, B., Fountzilas, G., Kalogeras, K.T., Kotoula, V., Papamichael, D., Laurent-Puig, P., et al.: Effects of kras, braf, nras, and pik3ca mutations on the efficacy of cetuximab plus chemotherapy in chemotherapy-refractory metastatic colorectal cancer: a retrospective consortium analysis. The Lancet oncology 11(8), 753–762(2010)

[31] Stelow, E.B., Mills, S.E.: Squamous cell carcinoma variants of the upper aerodigestive tract. American journal of clinical pathology 124 Suppl, 96–109(2005)

[32] Cho, N.-Y., Choi, M., Kim, B.-H., Cho, Y.-M., Moon, K.C., Kang, G.H.: Braf and kras mutations in prostatic adenocarcinoma. International journal of cancer 119, 1858–1862(2006). doi:10.1002/ijc.22071

[33] Park, J.Y., Kim, W.Y., Hwang, T.S., Lee, S.S., Kim, H., Han, H.S., Lim, S.D., Kim, W.S., Yoo, Y.B., Park, K.S.: Braf and ras mutations in follicular variants of papillary thyroid carcinoma. Endocrine pathology 24, 69–76(2013). doi:10.1007/s12022-013-9244-0

[34] Garcia-Rostan, G., Zhao, H., Camp, R.L., Pollan, M., Herrero, A., Pardo, J., Wu, R., Carcangiu, M.L., Costa, J., Tallini, G.: ras mutations are associated with aggressive tumor phenotypes and poor prognosis in thyroid cancer. Journal of clinical oncology : official journal of the American Society of Clinical Oncology 21, 3226–32 (2003). doi:10.1200/JCO.2003.10.13035.

[35] Silan, F., Gultekin, Y., Atik, S., Kilinc, D., Alan, C., Yildiz, F., Uludag, A., Ozdemir, O.: Combined point mutations in codon 12 and 13 of kras oncogene in prostate carcinomas. Molecular biology reports 39, 1595–1599(2012). doi:10.1007/s11033-011-0898-8

[36] Costa, A.M., Herrero, A., Fresno, M.F., Heymann, J., Alvarez, J.A., Cameselle-Teijeiro, J., García-Rostán, G.: Braf mutation associated with other genetic events identifies a subset of aggressive papillary thyroid carcinoma. Clinical endocrinology 68, 618–634(2008). doi:10.1111/j.1365-2265.2007.03077.x

[37] Paez, J.G., Jänne, P.A., Lee, J.C., Tracy, S., Greulich, H., Gabriel, S., Herman, P., Kaye, F.J., Lindeman, N., Boggon, T.J., Naoki, K., Sasaki, H., Fujii, Y., Eck, M.J., Sellers, W.R., Johnson, B.E., Meyerson, M.: Egfr mutations in lung cancer: correlation with clinical response to gefitinib therapy. Science (New York, N.Y.) 304, 1497–1500(2004). doi:10.1126/science.1099314

[38] Schmitt, M.W., Loeb, L.A., Salk, J.J.: The influence of subclonal resistance mutations on targeted cancer therapy. Nature reviews. Clinical oncology 13, 335–347(2016). doi:10.1038/nrclinonc.2015.175

[39] Zou, M., Baitei, E.Y., Alzahrani, A.S., BinHumaid, F.S., Alkhafaji, D., Al-Rijjal, R.A., Meyer, B.F., Shi, Y.: Concomitant ras, ret/ptc, or braf mutations in advanced stage of papillary thyroid carcinoma. Thyroid : official journal of the American Thyroid Association 24, 1256–1266(2014). doi:10.1089/thy.2013.0610

[40] Nagai, Y., Miyazawa, H., Huqun, Tanaka T., Udagawa, K., Kato, M., Fukuyama, S., Yokote, A., Kobayashi, K., Kanazawa, M., Hagiwara, K.: Genetic heterogeneity of the epidermal growth factor receptor in non-small cell lung cancer cell lines revealed by a rapid and sensitive detection system, the peptide nucleic acid-locked nucleic acid pcr clamp. Cancer research 65, 7276–7282(2005). doi:10.1158/0008-5472.CAN-05-0331

[41] Yamamoto, H., Oda, Y., Kawaguchi, K.-i., Nakamura, N., Takahira, T., Tamiya, S., Saito, T., Oshiro, Y., Ohta, M., Yao, T., Tsuneyoshi, M.: c-kit and pdgfra mutations in extragastrointestinal stromal tumor (gastrointestinal stromal tumor of the soft tissue). The American journal of surgical pathology 28, 479–488(2004)

[42] Jang, T.W., Oak, C.H., Chang, H.K., Suo, S.J., Jung, M.H.: Egfr and kras mutations in patients with adenocarcinoma of the lung. The Korean journal of internal medicine 24(1), 48 (2009)

[43] Boch, C., Kollmeier, J., Roth, A., Stephan-Falkenau, S., Misch, D., Grüning, W., Bauer, T.T., Mairinger, T.: The frequency of egfr and kras mutations in non-small cell lung cancer (nsclc): routine screening data for central europe from a cohort study. BMJ open 3(4), 002560 (2013)

[44] Vaqué, J.P., Martínez, N., Varela, I., Fernández, F., Mayorga, M., Derdak, S., Beltrán, S., Moreno, T., Almaraz, C., De las Heras G., et al.: Colorectal adenomas contain multiple somatic mutations that do not coincide with synchronous adenocarcinoma specimens. PLoS one 10(3), 0119946 (2015)

[45] Eskiocak, U., Kim, S.B., Ly, P., Roig, A.I., Biglione, S., Komurov, K., Cornelius, C., Wright, W.E., White, M.A., Shay, J.W.: Functional parsing of driver mutations in the colorectal cancer genome reveals numerous suppressors of anchorage-independent growth. Cancer research 71(13), 4359–4365(2011)

[46] Molina-Vila, M.A., Nabau-Moretó, N., Tornador, C., Sabnis, A.J., Rosell, R., Estivill, X., Bivona, T.G., Marino-Buslje, C.: Activating mutations cluster in the “molecular brake” regions of protein kinases and do not associate with conserved or catalytic residues. Human Mutation 35(3), 318–328(2014). doi:10.1002/humu.22493

[47] Hartmann, K., Wardelmann, E., Ma, Y., Merkelbach-Bruse, S., Preussner, L.M., Woolery, C., Baldus, S.E., Heinicke, T., Thiele, J., Buettner, R., Longley, B.J.: Novel germline mutation of kit associated with familial gastrointestinal stromal tumors and mastocytosis. Gastroenterology 129, 1042–1046(2005). doi:10.1053/j.gastro.2005.06.060

[48] Wang, Y.-Y., Zhou, G.-B., Yin, T., Chen, B., Shi, J.-Y., Liang, W.-X., Jin, X.-L., You, J.-H., Yang, G., Shen, Z.-X., Chen, J., Xiong, S.-M., Chen, G.-Q., Xu, F., Liu, Y.-W., Chen, Z., Chen, S.-J.: Aml1-eto and c-kit mutation/overexpression in t(8;21) leukemia: implication in stepwise leukemogenesis and response to gleevec. Proceedings of the National Academy of Sciences of the United States of America 102, 1104–1109(2005). doi:10.1073/pnas.0408831102

[49] Tamborini, E., Pricl, S., Negri, T., Lagonigro, M.S., Miselli, F., Greco, A., Gronchi, A., Casali, P.G., Ferrone, M., Fermeglia, M., Carbone, A., Pierotti, M.A., Pilotti, S.: Functional analyses and molecular modeling of two c-kit mutations responsible for imatinib secondary resistance in gist patients. Oncogene 25, 6140–6146(2006). doi:10.1038/sj.onc.1209639

[50] Norberg, S.T.: New phosphate langbeinites, k2mti(po4)3 (m = er, yb or y), and an alternative description of the langbeinite framework. Acta crystallographica. Section B, Structural science 58, 743–749(2002)

[51] Lee, J.C., Vivanco, I., Beroukhim, R., Huang, J.H.Y., Feng, W.L., DeBiasi, R.M., Yoshimoto, K., King, J.C., Nghiemphu, P., Yuza, Y., Xu, Q., Greulich, H., Thomas, R.K., Paez, J.G., Peck, T.C., Linhart, D.J., Glatt, K.A., Getz, G., Onofrio, R., Ziaugra, L., Levine, R.L., Gabriel, S., Kawaguchi, T., O’Neill, K., Khan, H., Liau, L.M., Nelson, S.F., Rao, P.N., Mischel, P., Pieper, R.O., Cloughesy, T., Leahy, D.J., Sellers, W.R., Sawyers, C.L., Meyerson, M., Mellinghoff, I.K.: Epidermal growth factor receptor activation in glioblastoma through novel missense mutations in the extracellular domain. PLoS medicine 3, 485 (2006). doi:10.1371/journal.pmed.0030485

[52] Kamata, T., Hussain, J., Giblett, S., Hayward, R., Marais, R., Pritchard, C.: Braf inactivation drives aneuploidy by deregulating craf. Cancer research 70, 8475–8486(2010). doi:10.1158/0008-5472.CAN-10-0603

[53] Olivier, M., Hollstein, M., Hainaut, P.: Tp53 mutations in human cancers: origins, consequences, and clinical use. Cold Spring Harbor perspectives in biology 2, 001008 (2010). doi:10.1101/cshperspect.a001008

[54] Zhang, Y., Coillie, S.V., Fang, J.-Y., Xu, J.: Gain of function of mutant p53: R282w on the peak? Oncogenesis 5, 196 (2016). doi:10.1038/oncsis.2016.8

[55] Rumi, E., Pietra, D., Ferretti, V., Klampfl, T., Harutyunyan, A.S., Milosevic, J.D., Them, N.C.C., Berg, T., Elena, C., Casetti, I.C., et al.: Jak2 or calr mutation status de?nes subtypes of essential thrombocythemia with substantially different clinical course and outcomes. Blood 123(10), 1544–1551(2013). doi:10.1182/blood-2013-11-539098

[56] Nangalia, J., Massie, C.E., Baxter, E.J., Nice, F.L., Gundem, G., Wedge, D.C., Avezov, E., Li, J., Kollmann, K., Kent, D.G., et al.: Somatic calr mutations in myeloproliferative neoplasms with nonmutated jak2. New England Journal of Medicine 369(25), 2391–2405(2013). doi:10.1056/nejmoa1312542

[57] Ha, J.-S., Kim, Y.-K.: Calreticulin exon 9 mutations in myeloproliferative neoplasms. Annals of Laboratory Medicine 35(1), 22 (2015). doi:10.3343/alm.2015.35.1.22

[58] Wang, M., Yang, C., Zhang, L., Schaar, D.G.: Molecular mutations and their cooccurrences in cytogenetically normal acute myeloid leukemia. Stem cells international 2017, 6962379 (2017). doi:10.1155/2017/6962379

[59] Wang, M., Yang, C., Zhang, L., Schaar, D.G.: Molecular mutations and their cooccurrences in cytogenetically normal acute myeloid leukemia. Stem cells international 2017 (2017)

[60] Dang, L., Yen, K., Attar, E.C.: Idh mutations in cancer and progress toward development of targeted therapeutics. Annals of oncology : official journal of the European Society for Medical Oncology 27, 599–608(2016). doi:10.1093/annonc/mdw013

[61] Duffy, M.J., Synnott, N.C., McGowan, P.M., Crown, J., O’Connor, D., Gallagher, W.M.: p53 as a target for the treatment of cancer. Cancer treatment reviews 40, 1153–1160(2014). doi:10.1016/j.ctrv.2014.10.004

[62] Pérez-Ramírez, C., Canadas-Garre, M., Molina, M.A., Faus-Dáder, M.J., Calleja-Hernández, M.A.: Pten and pi3k/akt in non-small-cell lung cancer. Pharmacogenomics 16, 1843–1862(2015). doi:10.2217/pgs.15.122

[63] Siena, S., Sartore-Bianchi, A., Di Nicolantonio, F., Balfour, J., Bardelli, A.: Biomarkers predicting clinical outcome of epidermal growth factor receptor-targeted therapy in metastatic colorectal cancer. Journal of the National Cancer Institute 101, 1308–1324(2009). doi:10.1093/jnci/djp280

[64] Paraiso, K.H.T., Xiang, Y., Rebecca, V.W., Abel, E.V., Chen, Y.A., Munko, A.C., Wood, E., Fedorenko, I.V., Sondak, V.K., Anderson, A.R.A., Ribas, A., Palma, M.D., Nathanson, K.L., Koomen, J.M., Messina, J.L., Smalley, K.S.M.: Pten loss confers braf inhibitor resistance to melanoma cells through the suppression of bim expression. Cancer research 71, 2750–2760(2011). doi:10.1158/0008-5472.CAN-10-2954

[65] Monticone, S., Hattangady, N.G., Nishimoto, K., Mantero, F., Rubin, B., Cicala, M.V., Pezzani, R., Auchus, R.J., Ghayee, H.K., Shibata, H., Kurihara, I., Williams, T.A., Giri, J.G., Bollag, R.J., Edwards, M.A., Isales, C.M., Rainey, W.E.: Effect of kcnj5 mutations on gene expression in aldosterone-producing adenomas and adrenocortical cells. The Journal of clinical endocrinology and metabolism 97, 1567–1572(2012). doi:10.1210/jc.2011-3132

[66] Rieke, D.T., Klinghammer, K., Keilholz, U.: Targeted therapy of head and neck cancer. Oncology research and treatment 39, 780–786(2016). doi:10.1159/000452432

[67] Leiserson, M.D.M., Vandin, F., Wu, H.-T., Dobson, J.R., Eldridge, J.V., Thomas, J.L., Papoutsaki, A., Kim, Y., Niu, B., McLellan, M., Lawrence, M.S., Gonzalez-Perez, A., Tamborero, D., Cheng, Y., Ryslik, G.A., Lopez-Bigas, N., Getz, G., Ding, L., Raphael, B.J.: Pan-cancer network analysis identifies combinations of rare somatic mutations across pathways and protein complexes. Nature genetics 47, 106–114(2015). doi:10.1038/ng.3168

[68] Lin, J., Gan, C.M., Zhang, X., Jones, S., Sjöblom, T., Wood, L.D., Parsons, D.W., Papadopoulos, N., Kinzler, K.W., Vogelstein, B., Parmigiani, G., Velculescu, V.E.: A multidimensional analysis of genes mutated in breast and colorectal cancers. Genome research 17, 1304–1318(2007). doi:10.1101/gr.6431107

[69] Dimitrakopoulos, C.M., Beerenwinkel, N.: Computational approaches for the identification of cancer genes and pathways. Wiley Interdisciplinary Reviews: Systems Biology and Medicine 9(1) (2017)

[70] Meador, C.B., Micheel, C.M., Levy, M.A., Lovly, C.M., Horn, L., Warner, J.L., Johnson, D.B., Zhao, Z., Anderson, I.A., Sosman, J.A., et al.: Beyond histology: translating tumor genotypes into clinically effective targeted therapies. Clinical Cancer Research 20(9), 2264–2275(2014)

